## Supplementary material for "Abnormal morphology and synaptogenic signaling in astrocytes following prenatal opioid exposure": Figure S1

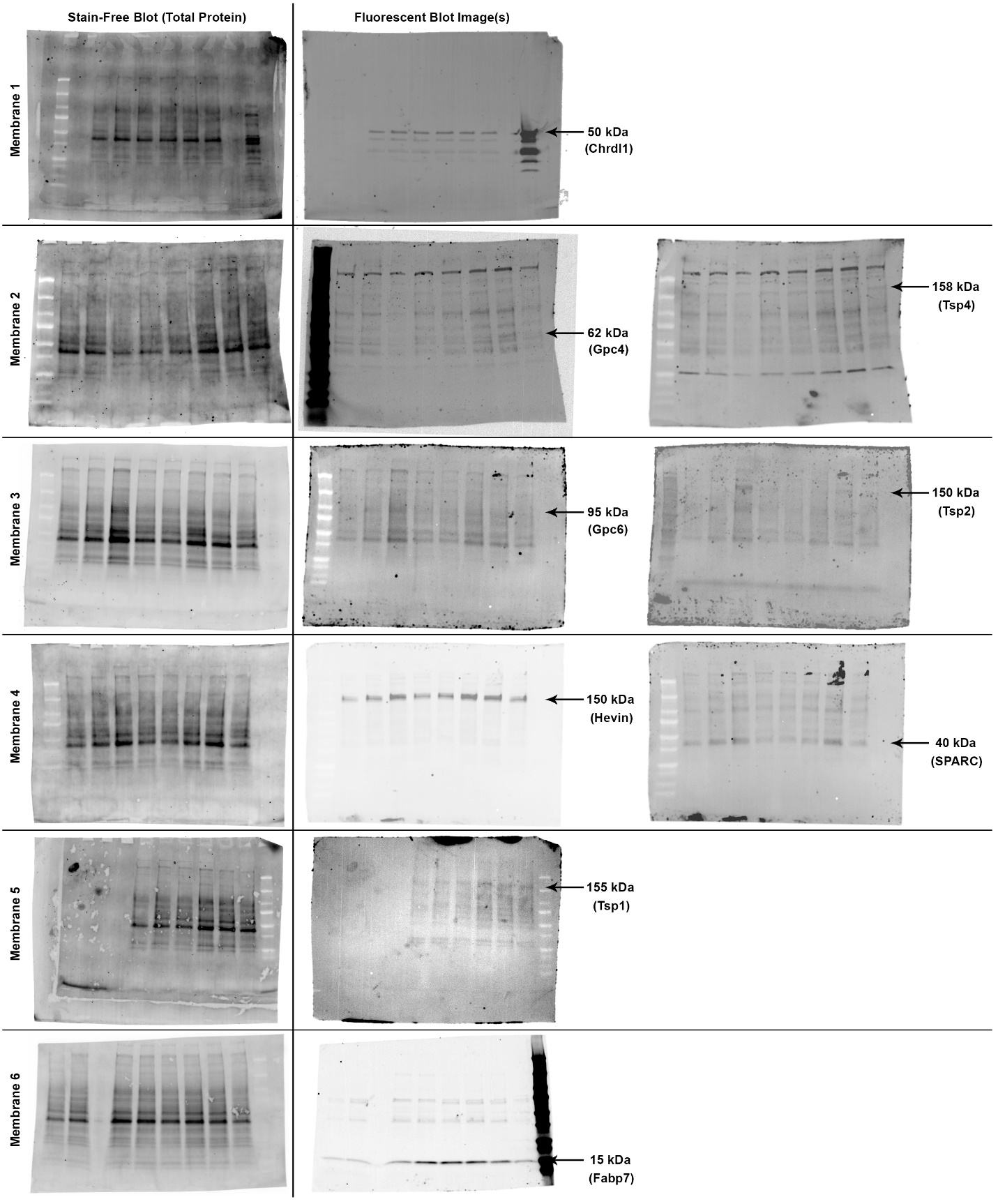

Uncropped Western blot membranes showing total protein content used for normalization along with the associated fluorescent blot images for the various primary antibodies probed.
